# Multi-omics insights into host-viral response and pathogenesis in Crimean-Congo Hemorrhagic Fever Viruses for novel therapeutic target

**DOI:** 10.1101/2020.12.10.419697

**Authors:** Ujjwal Neogi, Nazif Elaldi, Sofia Appelberg, Anoop T. Ambikan, Emma Kennedy, Stuart Dowall, Binnur K. Bagci, Soham Gupta, Jimmy E. Rodriguez, Sara Svensson-Akusjärvi, Vanessa M. Monteil, Ákos Végvári, Rui Benfeitas, Akhil C. Banerjea, Friedemann Weber, Roger Hewson, Ali Mirazimi

## Abstract

The pathogenesis and host-viral interactions of the Crimean–Congo hemorrhagic fever orthonairovirus (CCHFV) are convoluted and not well evaluated. Application of the multi-omics system biology approaches including biological network analysis in elucidating the complex host-viral response, allow for interrogating the viral pathogenesis. The present study aimed to fingerprint the system-level alterations during acute CCHFV-infection and the cellular immune responses during productive CCHFV-replication *in vitro*. We used system-wide network-based system biology analysis of peripheral blood mononuclear cells (PBMCs) from a longitudinal cohort of CCHF patients during the acute phase of infection and after one year of recovery (convalescent phase) and untargeted quantitative proteomics analysis of the most permissive CCHFV-infected Huh7 and SW13 cells. In the RNAseq analysis of the PBMCs, comparing the acute and convalescent-phase, we observed system-level host’s metabolic reprogramming towards central carbon and energy metabolism (CCEM) with distinct upregulation of oxidative phosphorylation (OXPHOS) during CCHFV-infection. Upon application of network-based system biology methods, negative coordination of the biological signaling systems like FOXO/Notch axis and Akt/mTOR/HIF-1 signaling with metabolic pathways during CCHFV-infection were observed. The temporal quantitative proteomics in Huh7 showed a dynamic change in the CCEM over time and was in agreement with the cross-sectional proteomics in SW13 cells. By blocking the two key CCEM pathways, glycolysis and glutaminolysis, viral replication was inhibited *in vitro*. Activation of key interferon stimulating genes during infection suggested the role of type I and II interferon-mediated antiviral mechanisms both at system-level and during progressive replication.

**Significance Statement:** A combination of multi-modal systems-wide host-immune response and *in vitro* temporal analysis identified molecular re-arrangement in CCEM and fingerprinting the interferon-mediated antiviral mechanism during CCHFV-infection. Using the newly gained insights, we then modulated the key pathways of CCEM by drugs and inhibited the productive CCHFV-replication in *in vitro* infection models. Our study thus provides a comprehensive, system-level picture of the regulation of cellular and metabolic pathways during productive CCHFV-infection for the first time that aids in identifying novel therapeutic targets and treatment strategies.

## Introduction

Crimean–Congo hemorrhagic fever orthonairovirus (CCHFV), a negative-sense RNA virus belonging to the *Nairoviridae* family, is a major emerging pathogen with an increasing number of outbreaks all over the world. Causing a mild to severe viral hemorrhagic fever (CCHF; Crimean– Congo hemorrhagic fever) poses a substantial threat to public health due to its high mortality rate in humans (3–40%), modes of transmission (tick-to-human/animal, animal-to-human, and human-to-human) and geographical distribution. CCHF is endemic in almost 30 countries in sub-Saharan Africa, South-Eastern Europe, the Middle East, and Central Asia (1, 2). The ixodid ticks, especially those of the genus *Hyalomma*, are both a vector and a reservoir for CCHFV and are highly ubiquitous with their presence in more than 40 countries (3). In recent years, CCHFV outbreaks have become more frequent and expanded to new geographical areas. This has been attributed to climate change and the spread of infected ticks by birds and the livestock trade. The presence of the CCHFV tick vector in Portugal, Spain, Germany, and even Sweden (4) and England (5) highlights the need for stricter surveillance due to the possibility of a future intrusion (6). Turkey has reported the highest number of laboratory-confirmed CCHF cases and is one of the worst affected countries in the world (7). Since the first identification in 2002 up till the end of 2019, a total of 11,780 confirmed CCHF cases have been reported with a case-fatality rate of 4.7% (unpublished data by the Turkish Ministry of Health). On average, there were nearly 500 cases every year, reported mainly during the summer months May-July (8).

Because of the sporadic nature of CCHF outbreaks in humans in the endemic regions, a lack of infrastructure, and the absence of systematic studies, little is known about the pathogenesis and host-virus interactions during the acute phase of CCHF disease and associated sequelae after recovery. An in-depth understanding of host responses to CCHFV is necessary to design better therapeutic and containment strategies for CCHF. Systems biology studies using -omics approaches on patient material and infected cells can elucidate potential mechanisms of the host immune response, disease pathogenesis, and potential host response that can distinguishes disease severity as reported recently in 16 viruses including severe acute respiratory syndrome coronavirus 2 (SARS-CoV-2), Chikungunya, Zika, Ebola, Influenza viruses (9-11). However, no data was reported for CCHFV infection.

Here, we have applied global blood transcriptomics in longitudinal samples collected during both the acute phase of CCHFV-infection and the convalescent phase (nearly after a year of recovery) to measure the system-wide changes during the CCHFV-infection in patients from Turkey. We also performed temporal quantitative proteomics analysis to understand the cellular alterations during the productive CCHFV-infection in two different cell lines, human adrenal carcinoma cell line, SW13 and human liver cell line Huh7 that were reported to be the most permissive cell lines for CCHFV (12). Using the newly gained insights, we then modulated the key pathways by drugs to halt the productive CCHFV-replication in *in vitro* infection models. Our study thus provides a comprehensive, system-level picture of the regulation of cellular and metabolic pathways during productive CCHFV-infection that can aid to identify novel therapeutic targets and strategies.

## Results

### Samples and clinical data

In this study, 18 samples were collected during the acute phase of the disease with a median time of 4 days (range 1-6 days) after the onset of symptoms. By using severity grading scores (SGS), 33% (6/18) patients were grouped into severity group 1 (SG-1), 61% (11/18) patients into severity group 2 (SG-2) and, 6% (1/18) patients into severity group 3 (SG-3). The median age of the patients was 49 years (range: 18–79), and 12 (66.7%) of the patients were male. A 79-year-old male patient in SG-3 died on the third day of hospitalization. The case-fatality rate (CFR) for the cohort was 5.6%. Follow-up samples were collected from 12 individuals after a median duration of 54 weeks (range: 46-57 weeks). The CCHF patient characteristics are summarized individually in Table S1.

### System-level metabolic reprogramming during the acute phase of CCHFV infection

Due to the natural heterogeneity in human cohorts, we used longitudinal samples from 12 patients (SG-1: n=5; SG-2: n= 7) to perform differential expression analyses for each infected patient between the time of infection and approx. 1-year post-recovery (Range: 46–57 weeks). The differential gene expression (DGE) profile for the acute phase compared to the recovered phase in all patients showed an upregulation of 2891 genes and a downregulation of 2738 genes (Fig 1A and Dataset S1). Next, we used the functional analysis using a consensus scoring approach based on multiple gene set analysis (GSA) runs by incorporating the directionality of gene abundance using R/Bioconductor package PIANO (13) for KEGG-2021. Using the group-specific consensus scores (acute vs. recovered) and directionality classes, we identified distinct upregulation (adj. p<0.05) of metabolic pathways such as one carbon pool by folate, oxidative phosphorylation (OXPHOS), glycolysis, N-glycan biosynthesis, and antiviral pathways like the NOD-like receptor signaling pathway (Fig. 1B and Dataset S2). However, the pathways related to the down-regulated genes were mainly antiviral defense mechanism-associated pathways including innate immune responses like Th1, Th2 and Th17 cell differentiation, the NF-kB pathways, chemokine signaling pathway, etc. (Fig. 1B). Additionally, given that most of the metabolic pathways were upregulated, we used the DGE results of acute-vs-recovered to identify reporter metabolites. Reporter metabolites are metabolites around which most of the transcriptional changes occur (14) thus being indicative of gene-level altered regulation of metabolism. The analysis identified 37 significantly upregulated reporter metabolites (adj. p<0.1), that were part of OXPHOS, TCA-cycle, nucleotide metabolism, N-glycan metabolism, and amino acid-related pathways (Fig 1C). To specifically investigate the genes that were significantly associated with disease severity during the acute phase, the samples were grouped into either SG-1 or SG-2 and 3 combined. There were 12 genes (*ERG, PROM1, HP, HBD, AHSP, CTSG, PPARG, TIMP4, SMIM10, RNASE1, VSIG4, CMBL, MT1G*) that were significantly upregulated in patients in the SG-2 and SG-3 combination group compared to SG-1 (Fig. S1A) However, no obvious links between these genes were noted and no apparent clustering was observed (Fig S1B). This was further supported by serum secretome analysis using the 22 soluble cytokine and chemokine markers by Luminex assay on samples collected during the acute phase of the disease from SG-1 (n=6) and SG-2 (n=11). Of the 22 markers used for analysis, only interleukin 8 (IL-8) and Granulocyte-macrophage colony-stimulating factor (GM-CSF) was shown borderline significance between SG-1 and SG-2 (Fig S2). However, when we compared the acute phase with the recovered phase in SG-1, and SG-2 separately, there was a distinct DGE profile. In SG-1 the differentially expressed genes were significantly fewer (adj. p<0.05; n=1617, upregulated: 954 and downregulated: 663) compared to those in SG-2 (n=4256, upregulated: 2182 and downregulated: 2074) (Fig. 1D and 1E). There were 1451 overlapping genes between SG-1 and SG-2 that were differentially upregulated (n=882) and downregulated (n=569). Using gene ontology (GO) analysis after removal of the redundant terms using REVIGO (15), the majority of the genes from the top two GO terms that were significantly upregulated were part of the IFN-I signaling pathway (GO:0060337) and the regulation of viral genome replication (GO:0045069) (Fig. 1F). This indicates that the disease severity had a significant effect on gene expression of the interferon signaling pathway profiling during the acute phase, whereas it was comparable when they recovered.

**Figure 1.**
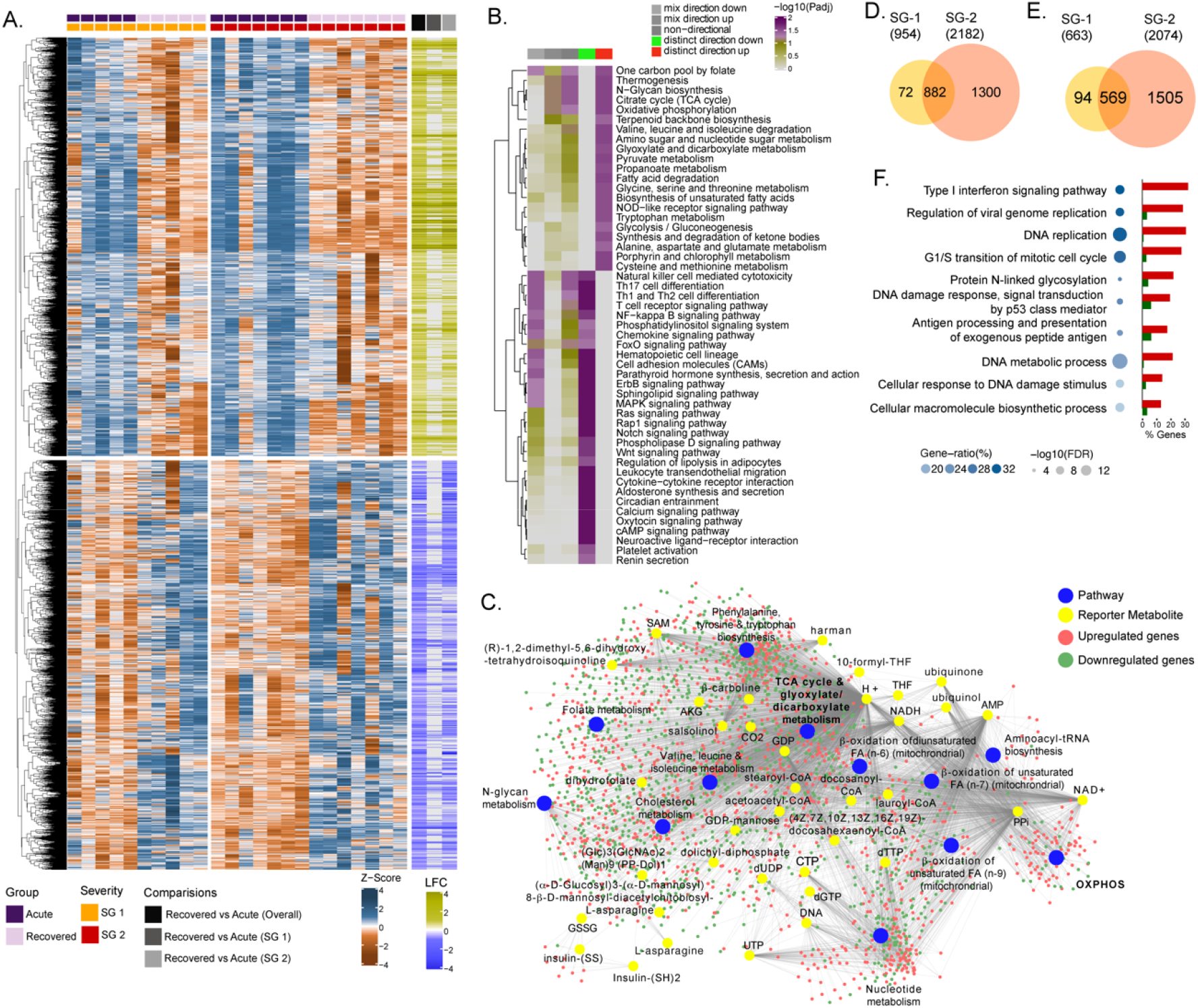
Differential gene expression and pathway analysis between acute and recovery phases. **(A)** Heatmap of Z-score transformed expression values of significantly regulated genes in the pair-wise comparisons namely recovered vs. acute (overall), recovered vs. acute (SG-1), recovered vs. acute (SG-2). The columns represent the patient samples and their corresponding severity groups at different time points. The rows represent genes that are hierarchically clustered based on Euclidean distance. **(B)** Pathways found to be significantly regulated (adj. p<0.05) by genes expressed at the acute infection phase compared to recovered phase. The heatmap visualizes negative log scaled adjusted p-values of different directionality classes. Non-directional p-values are generated based on gene level statistics alone without considering the expression direction. The mixed-directional p-values are calculated using subset of gene level statistics of up and down regulated genes respectively for mixed-directional up and down. Distinct directional up and distinct directional down p-values are calculated from gene statistics with expression direction (**C)** Network visualization of significant reporter metabolites (adj. p<0.1) and reporter subsystems identified in acute compared to recovered. Yellow node denotes reporter metabolite and blue node denotes reporter subsystems. Light red and green colored nodes represent upregulated and downregulated genes respectively. Each edge in the network denotes association of genes with reporter metabolites and subsystems based on human genome scale metabolic model. **(D)** Venn diagram of significantly up-regulated genes in recovered vs acute (SG-1) and recovered vs acute (SG-2) phases **(E)** Venn diagram of significantly down-regulated genes in recovered vs. acute (SG-1) and recovered vs. acute (SG-2) phases. **(F)** Gene ontology (GO, biological process) enrichment analysis results of commonly regulated genes (882 upregulated and 569 down-regulated) from (D) and (E). The color gradient and bubble size correspond to the gene ratio of each GO term and the adjusted P value of the enrichment test, respectively. The adjacent bar graph represents the percentage of genes upregulated or downregulated in each GO term.

### Distinct interferon signaling-related pathways in CCHFV-infection

To identify the CCHFV-induced changes in the interferon-related signaling pathways, we used our previously curated datasets for genes (n=205) associated with the interferon response (16). The majority of the genes of the interferon signaling pathways were upregulated (36%, 73/205, adj p<0.05) while 11% (22/205) were downregulated (Fig 2A). Of the IFN-regulated genes, IFI27 (ISG12) showed the most robust upregulation (Fig. 2B). This was further supported by RNAscope™ analysis targeting the IFI27 transcript in the SW13 cell line infected with CCHFV strain IbAr10200 (Fig. 2C). Apart from this ISG20, ISG15, Mx1, Mx2 and several other ISGs showed upregulation in the acute phase (Dataset S1). Given that the interferon signaling pathways have a role in disease severity, we next performed association between the patient viral load and genes in the interferon signaling-related pathways. We identified six genes (*TRIM25, IFI35, EIF2AK2, USP18, IFI6 and BST2*) that were negatively associated with the cycle threshold (CT) value of RT-PCR (adj p<0.05 and R>-0.82, Fig. 2D), suggesting a higher viral load was associated with an increased expression of these ISGs. Overall, the gene expression data indicate that the CCHFV infection regulates IFN responses and patients with a successful disease outcome showed stimulation of several ISGs during the acute phase of infection.

**Figure 2.**
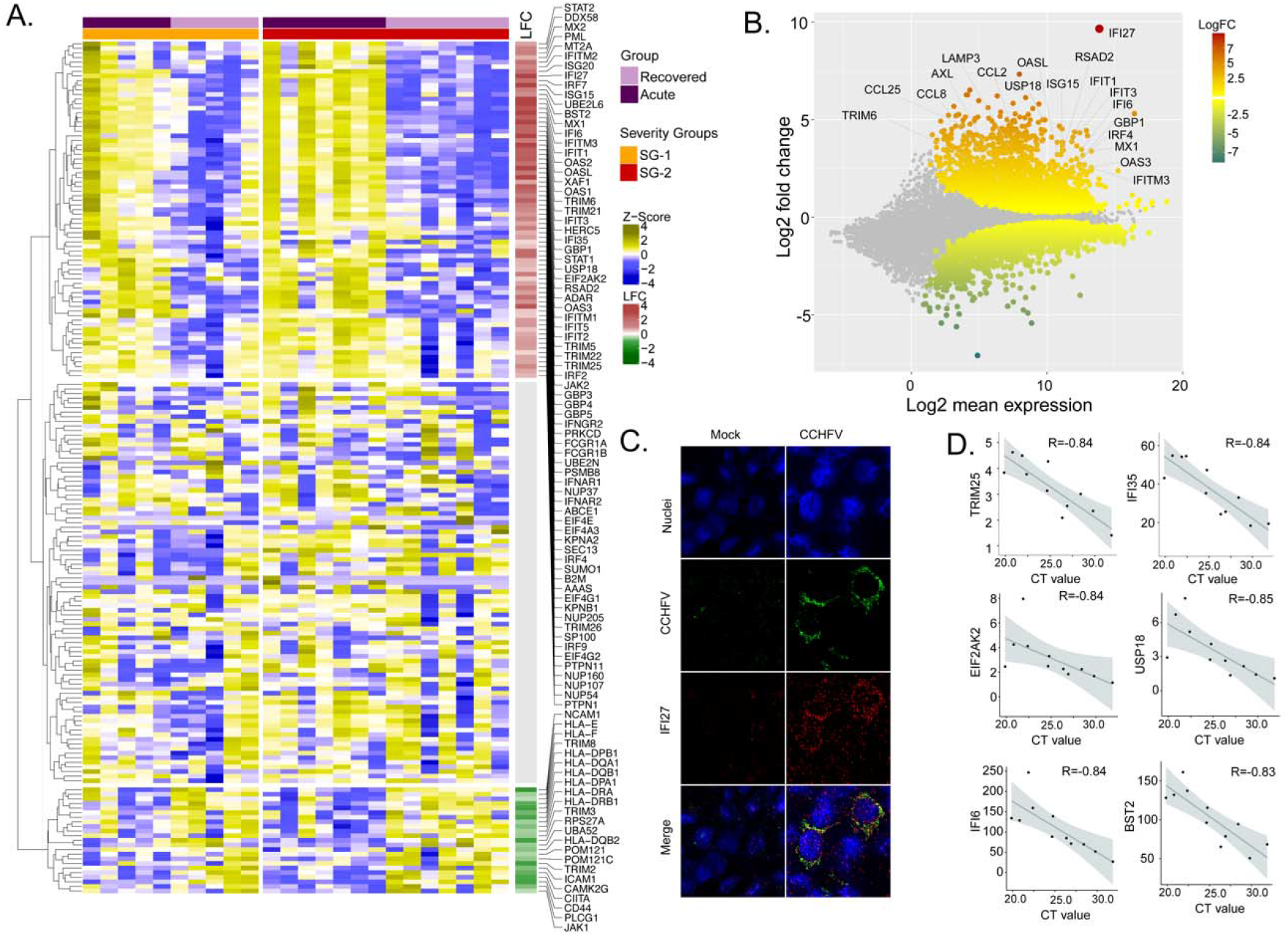
Differentially expressed genes in interferon (IFN) signaling pathways. **(A)** Heatmap visualizing the expression pattern of IFN-signaling genes (including ISGs) that are significantly different between the recovered and acute phases. The columns represent the patient samples and their corresponding severity groups at different time points. The rows represent genes hierarchically clustered based on Euclidean distance. The row annotation indicates the differential gene expression of genes (DGE) and presence or absence of genes in the selected GO terms. The names of genes involved in the cellular response to IFN-γ (GO:0071346) and the IFN-I signaling (GO:0060337) are printed. **(B)** MA-plot of differentially regulated genes between the recovered and acute phases. ISGs are marked. **(C)** RNAscope analysis targeting IFI27 genes in infected and non-infected cells. **(D)** Network type visualization of genes belonging to GO:0071346 and GO:0060337. The edges indicate the presence of the corresponding gene in the gene-ontology term. The node size and color gradient correspond to the adjusted P value of differential expression analysis and the log2 fold change, respectively. (**E**) Spearman correlation between viral load and IFN signaling genes (adj p<0.05).

### Network analysis identified the central role of central carbon and energy metabolism (CCEM) in the regulation of signaling pathways

To further deepen our understanding of the cellular regulation of acute CCHFV-infection at the molecular level from a systems perspective, we employed a weighted gene co-expression network analysis at the transcriptomic level. Based on the network analysis of pairwise gene co-expression (adj. p<0.001, Spearman ρ>0.84), we identified a set of seven communities of strongly interconnected genes (Fig. 3A). Next, we ranked all the communities based on their centrality to identify the sets of genes with the highest coordinated expression changes that were predicted to robustly influence network behavior. The functional enrichment analysis of the central community (c1) of the transcriptomics is associated (adj. p<0.05) mainly with alterations in pyruvate metabolism, TCA-cycle and to a smaller extent to glycolysis and gluconeogenesis (adj. p<0.2) (Fig 3A and Fig S3). Further, we observed (Fig 3B and S4) a high number of negative correlations between community (c1) and those associated with Notch, mechanistic target of rapamycin (mTOR) and Forkhead box protein O (FoxO) signaling (c5) and hypoxia inducing factor-1 (HIF-1) signaling (c7). Interestingly, the OXPHOS-associated community (c3) also tends to be negatively correlated with those involved in Notch/mTOR/FoxO signaling (c5) and HIF-1 signaling (c7). These patterns are also observed among the top 10% of most central genes in each community, suggesting key opposite differences not only at a global community level but also in key genes in each community (Fig 3B). At a pathway level, we indeed observed antagonistic trends between the above-mentioned pathways (Fig. 3C). Our functional and network community analyses in the patient transcriptomics identified the coordination of biological signaling systems like FoxO, Notch and mTOR/HIF-1 signaling with metabolic pathways of CCEM during CCHFV-infection.

**Figure 3.**
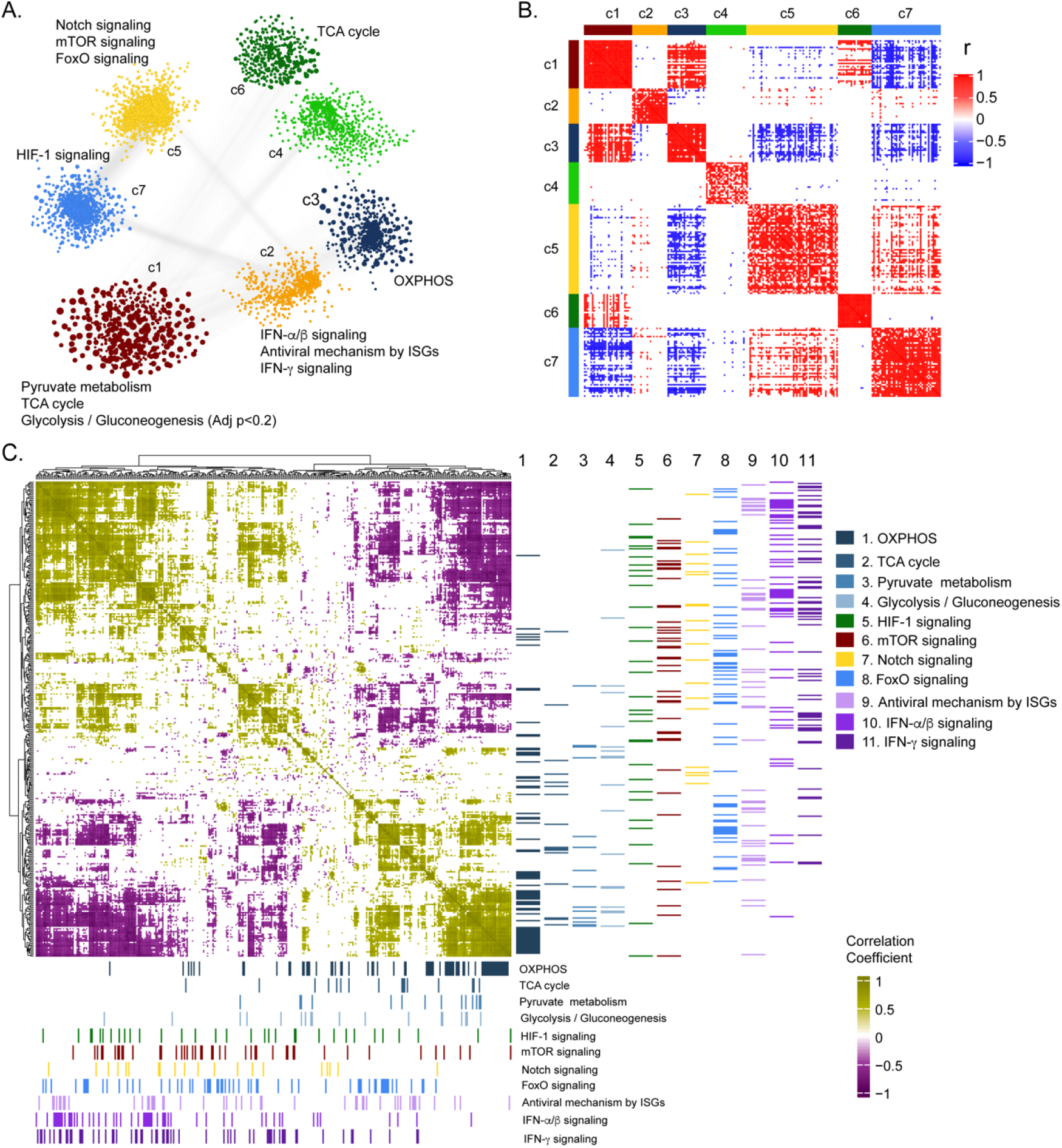
Weighted co-expression network analysis. **(A)** Network visualization of seven gene co-expression communities identified. Nodes and node size represent genes and their centrality (degree) respectively and edges represent significant spearman correlation (adj p<0.001 and R>84). Key significantly regulated pathways (padj <0.05) in each community are labelled. **(B)** Heatmap of correlations among top 5% central genes in each community. Column and row annotation denotes corresponding communities. **(C)** Heatmap of significant correlation (adj. p<0.05) between key metabolic and signaling pathways mentioned in (A). Column and row annotation denotes corresponding pathways.

### Quantitative proteomics analysis identified modulation of key metabolic processes and signaling pathways during productive replication *in vitro*

Our longitudinal transcriptomics analysis of CCHF patient samples revealed alternations in the several key metabolic processes and signaling pathways during the acute phase of infection at a system level. As CCHFV fails to infect the peripheral blood mononuclear cells (PBMCs) (17), to understand the global changes in the cellular response during productive CCHFV-infection, we infected Huh7 and SW13 cells with CCHFV, which are the common cell lines used in pathogenesis studies and considered highly permissive for CCHFV (12). To allow multiple rounds of infection we used a multiplicity of infection (MOI) of 1 and used a time-course proteomic experiment for 24 and 48hpi using single batch TMT-labelling based mass-spectrometric analysis to avoid batch effects, inflated false-positive results and minimize the typical missing values issue (18). Due to the higher cell death the proteomics analysis could not be performed in SW13 48hpi. In the UMAP clustering of the proteome data, we observed a clear separation between the mock and virus-infected cells in both the cell lines (Fig. 4A). At 24hpi and 48hpi a substantial amount of CCHFV proteins, N, M, and L protein were detected (Fig. 4B). The immune fluorescence analysis targeting N-protein of CCHFV infected Huh7 cells at 24hpi with 1 MOI is shown in Fig 4C. The differential protein analysis (DPA) identified 3205 and 3070 proteins upregulated and 2926 and 3279 proteins downregulated in the infected samples at 24hpi and 48hpi in Huh7 cells and 2217 upregulated and 1705 downregulated in SW13 cells respectively compared to the mock (adj. p<0.05). The consensus scoring based protein set analysis (PSA) using PIANO on the DPA at 24hpi and 48hpi in Huh7 and 24hpi in SW13 identified 68 pathways to be dysregulated in at least one of the comparisons. We observed downregulation (adj. p<0.05) of the glycolysis/gluconeogenesis, purine metabolism, PI3K-Akt, and HIF-1 signaling pathways in both Huh7 and SW13 cell lines at 24hpi (Fig 4D) indicating CCHFV utilized these pathways during productive replication at an early phase. These pathways are known to have feedback mechanisms (19, 20) to maintain cellular homeostasis, which is consistent with the observation that at 48hpi (in Huh7 cells) the pathways were not significantly dysregulated. The pathways like TCA-cycle and insulin secretion showed opposite trends in the cell lines indicating cell type-specific differential regulation of metabolic and signaling pathways during CCHFV replication. In time-series analysis in Huh7 cells, oxidative phosphorylation (OXPHOS) pathway was upregulated during CCHFV infection in a temporal manner indicating shift in metabolic processes towards OXPHOS during productive replication of the virus. The other pathways that also showed distinct temporal upregulation during CCHFV infection in vitro were N-glycan biosynthesis and cytokine-cytokine receptor interactions. In turn, pathways like FoxO signaling, T-cell receptor signaling pathways, Th1 and Th2 cell differentiation, and NK cell-mediated cytotoxicity were downregulated and upregulated of Notch signaling in Huh7 24hpi but not at 48hpi indicated the role of these pathways at the early stage of infection. A severe metabolic rearrangement occurred in SW13 cells at 24hpi towards central carbon and energy metabolism and amino acid metabolism as the pathways like pyruvate metabolism, glycine, serine and threonine metabolism, tryptophan metabolism etc. were downregulated (Fig 4D). We also performed quantitative proteomics analysis of the Huh7 cells with 4 MOI infections at 24hpi and observed similar alterations in the pathways (Fig S5). Next, we performed gene and protein set enrichment analysis (GSEA and PSEA) in Enricher and compared Huh7 and SW13, 24hpi and patients RNAseq data and observed the key common dysregulated pathways were TCA cycle, HIF-1 and FoxO signaling pathways (Fig 4E). Glycolysis and OXPHOS are molecular interconversion systems, where the end product of the glycolysis is fueling OXPHOS through the TCA cycle which normally is the primary energy source and major pathways of CCEM. Glutaminolysis is an alternative pathway for mitochondrial energy production through OXPHOS under altered metabolic conditions (21, 22). Therefore, we blocked glycolysis and glutaminolysis, in SW13 and Huh7 cells, using 2-deoxy-D-glucose (2-DG) (5mM) and 6-diazo-5-oxo-L-norleucine (DON) (50μm) respectively (Fig. 4F) following infection. Infectivity of CCHFV, quantified as relative CCHFV *L-gene* levels in cells lysates, showed significant decrease in 2-DG treated cells in both SW13 and Huh7 (p=0.003 and p=0.028 respectively). While in the DON treated cells a significant decrease was observed in SW13 cell (p<0.001) and an inhibitory trend in Huh7 (p=0.162) (Fig 4G). These data indicate that alteration in the CCEM affects CCHFV replication despite the cell-specific differences.

**Figure 4.**
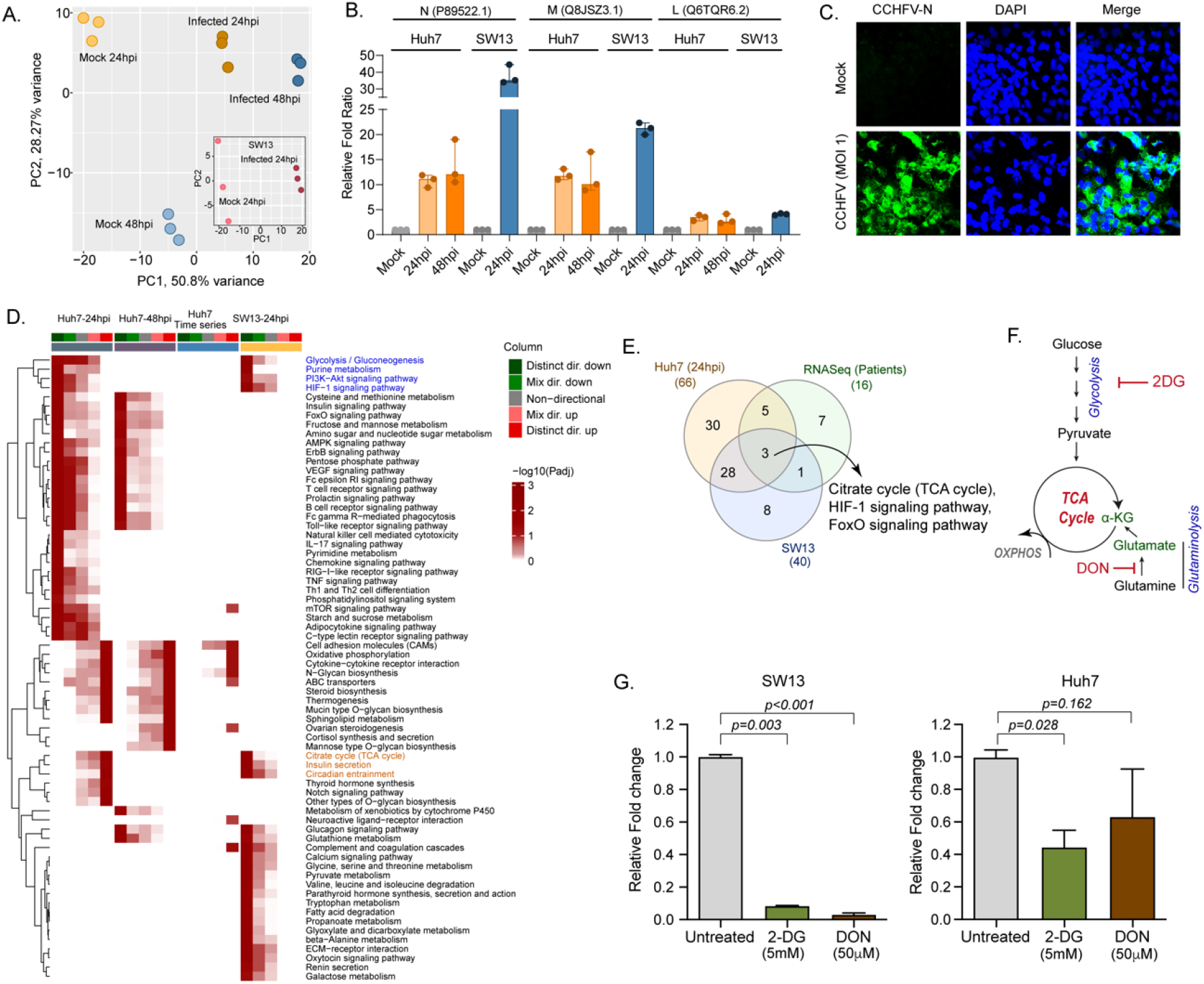
LC-MS/MS based quantitative proteomics analysis in CCHFV-infected Huh7 and SW13 cells. **(A)** Principal component analysis of proteomics samples of Huh7 cells and SW13 (inset). **(B)** Identification of the CCHFV N (UniProtKB P89522.1), M (UniProtKB Q8JSZ3.1) and L (UniProtKB Q6TQR6.2) protein in the quantitative proteomics analysis. **(C)** Immunofluorescence staining of the CCHFV nucleoprotein to assess the infectivity. **(D)** Significantly regulated pathways (adj p<0.05) in any of the pair-wise proteomics analysis in Huh7 and SW13 cells. The heatmap visualizes negative log scaled adjusted p-values of different directionality classes. Non-directional p-values are generated based on gene level statistics alone without considering the expression direction. The mixed-directional p-values are calculated using subset of gene level statistics of up and down regulated genes respectively for mixed-directional up and down. Distinct directional up and distinct directional down p-values are calculated from gene statistics with expression direction. First column annotation represent directionality of pathways and second column annotation denotes corresponding differential expression analysis. **(E)** Venn diagram showing common dysregulated pathways in patients transcriptomics and cell line proteomics. (**F**) Schematic diagram of the glycolysis and glutaminolysis and targeted drugs. (**G**) Metabolic control of viral replication *in vitro*. Fold change of the CCHFV *L-gene* following infection and treatment of 2-DG and DON at indicated concentrations compared to untreated in SW13 cells and Huh7 cells. A two-tailed paired Student *t* test was performed, and p values are mentioned.

### Temporal dynamics of interferon response *in vitro*

The temporal changes in the interferome (cluster of interferon genes) are represented as a heat-map in Fig. 5A and the log2fold change of the significantly altered protein levels at 24hpi and 48hpi are represented as volcano plots in Fig. 5B. Several ISGs, such as Mx1, Mx2 IFIT1, ISG15, ISG20, and IFI6, were transcriptionally upregulated in the acute phase in patient samples (Supplementary Data File 1), were also significantly elevated in proteomics of infected Huh7 cells by 48hpi (Fig. 5C). To determine that the observed induction of ISGs is due to the CCHFV-infection itself and not caused by presence of any residual interferon in the virus-containing supernatant, we performed infection using UV-inactivated virus supernatant. As shown in the immunoblots in Fig. 5D, significant increase in expression of several ISGs namely RIG-I, IFIT1, ISG15 and a noticeable increase in Mx1, Mx2 and ISG20 proteins were observed in CCHFV-infected cells and not in UV inactivated virus supernatant, confirming that CCHFV-infection induces the expression of these ISGs.

**Figure 5.**
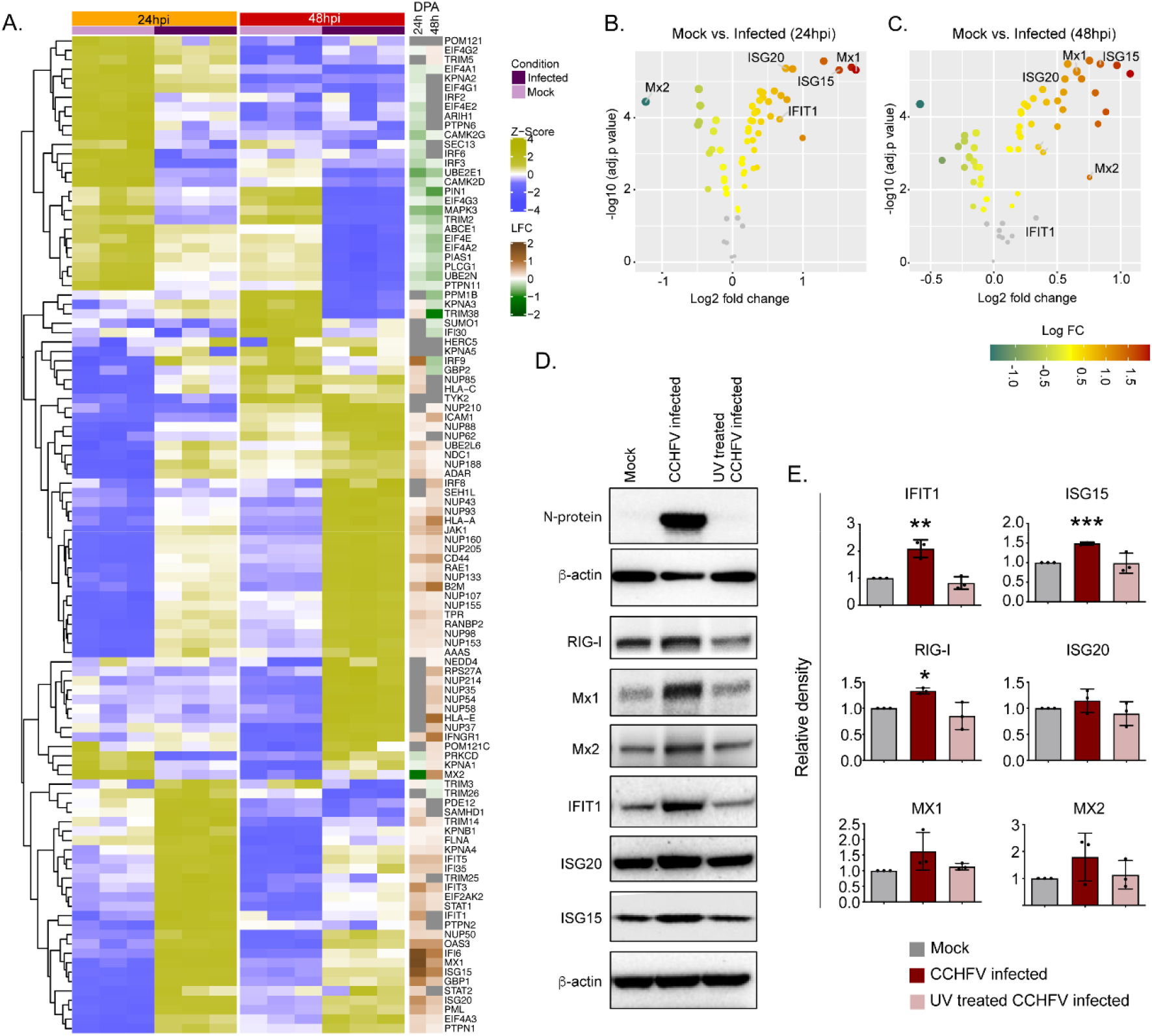
Temporal dynamics of interferon stimulating genes (ISGs). **(A)** Heatmap of Z-score transformed expression values of proteins belonging to the cellular response to IFN signaling pathways in Mock-infected and CCHFV-infected Huh7 cells at 24hpi and 48hpi as identified in proteomics. The log-2-fold change in the genes corresponding to the indicated proteins identified in our patient transcriptomics data (recovered vs acute) is shown under the column name RNASeq. **(B and C)** Volcano plot of ISGs visualizing the expression status of Mock-infected and CCHFV-Infected samples at **(B)** 24hpi and **(C)** 48hpi. The size and colour gradients of the dots correspond to the adjusted P values of differential expression analysis and the log2 fold change, respectively. **(D)** Representative western blots illustrating the expression of the indicated ISGs in Mock-infected, CCHFV-infected, and UV-inactivated CCHFV-infected Huh7 cells at 48hpi. ISG20 antibody gave a specific band at approx. 40kDa without any non-specific band in the membrane that was cut at 50kDa in the top. (**E**) The densitometric intensity of the bands were quantified using Fiji (ImageJ) software. The intensity of the individual bands was first normalized to the respective β-actin loading control and further relative normalization with respect to the mock-infected control was done. The bars are represented as means±SD of three independent experiments. A two-tailed paired Student *t* test was performed, and p values are represented as *P<0.05, **P<0.01 and ***P<0.001.

## Discussion

In our study, using the system level genome-wide transcriptomic analysis of a longitudinal patient cohort, temporal quantitative proteomics from *in vitro* infection assays in Huh7 cells, cross-sectional quantitative proteomics analysis in SW13 cells and *in vitro* inhibition of CCHFV replication following the blocking the glycolysis and glutaminolysis, we showed that during CCHFV-infection there is metabolic reprogramming of host cells towards central carbon and energy metabolism and that this plays a major role in viral replication despite the existence of cell-type-specific differences. Upregulation of OXPHOS was a unique feature of CCHFV-infection, at both the system-level blood transcriptomics and cellular proteomics during productive infection in Huh7. By applying network-based system biology methods, we identified the negative co-ordination of the biological signaling systems like FoxO/Notch axis and mTOR/HIF-1 signaling along with metabolic pathways of CCEM during CCHFV-infection at the system level. Blocking the two key CCEM pathways, glycolysis and glutaminolysis, controlled viral replication *in vitro*. Moreover, IFN-I mediated antiviral mechanisms were also activated with elevated key antiviral ISGs (ISG12, ISG15, ISG20), and MXs (Mx1 and Mx2).

Viruses exploit the host metabolic machinery to meet their biosynthetic demands. This reliance is further highlighted by observed variations in the cell-specific viral replications and production leading to changes in host metabolism (23). The changes in the energy metabolism can therefore be seen as an evolving property of the combined host-virus metabolic system and could be related to changes in host cellular demands arising from viral production (24). Our system-level transcriptomics data on patient material and *in vitro* cell culture assays indicated a transient dysregulation of key metabolic processes of the CCEM, like OXPHOS, glycolysis, and TCA-cycle in CCHFV-infection. These pathways are also known to promote replication of several other RNA viruses including human immunodeficiency virus type 1 (HIV-1), rubella virus, dengue virus (DENV), rhinovirus, hepatitis C virus (HCV), influenza virus etc (25, 26). Blocking glycolysis and glutaminolysis, that fuel OXPHOS, resulted in severe suppression of CCHFV-replication suggesting the need for these pathways for efficient viral replication. Our system biology analysis further indicated the coordinating role of the metabolic pathways of CCEM with biological signaling systems like Notch/FoxO axis and mTOR/HIF-1 signaling during the CCHFV-infection. It is known that these biological systems regulate energy metabolism. Notch signaling plays an essential role to maintain the cellular energy homeostasis via regulation of HIF-1 and PI3K/AKT signaling that is known to induce glycolysis (27).

On the other hand, FoxO signaling regulates cell proliferation by modulating energy metabolism and gluconeogenesis (28). The coordinated role of these transcriptional regulators (HIF-1α, FoxO, mTOR, and Notch1) modulates OXPHOS and mitochondrial biogenesis (29). Notch signaling has also been known to facilitate viral infectivity of RNA viruses including influenza virus, respiratory syncytial virus (RSV), HCV etc. (30) and have regulatory roles in inflammation (31). Our study is concordant with an earlier study that reported the downregulation of the Notch signaling in CCHFV-infection at the transcript level (32). However, our study also pointed out that during productive infection in Huh7 cells, Notch signaling was upregulated at 24hpi but not at 48hpi, indicating a role in the early stage of viral replication. Silencing of the Notch1 reported increasing toll like receptor 4 (TLR4) triggered proinflammatory cytokines (33) which is common during acute CCHFV-infection (34).

Several viruses encode proteins such as Ebola virus (EBOV) glycoprotein, the Dengue virus (DENV) nonstructural protein 1 (NS1), *etc*., that are known to activate TLR4 (35). On the other hand, at the first encounter with the pathogen, PI3Ks negatively regulate TLRs including TLR4 signaling (36). Of note, in our proteomics data during productive infection, we observed downregulation of PI3K/Akt signaling at 24hpi but not at 48hpi. Though there was no distinct downregulation of the whole pathway, in our patient system-level transcriptomics data, we also noted significant downregulation of genes belonging to the PI3K/Akt pathway during acute CCHFV-infection. Moreover, apart from PI3K/Akt, mTOR and HIF-1 signaling were also downregulated at 24hpi indicating modulation of PI3K/mTOR/HIF-1 axis by CCHFV for its replication. In our previous study we have shown that exogenous nitric oxide that is known to regulate the HIF-1 via the Akt/mTOR pathway under normoxic conditions (37), inhibited CCHFV *in vitro* (38). Interestingly, an *in vitro* study in another *Bunyavirus*, Rift Valley fever virus (RVFV), identified the inhibition of the PI3K/Akt pathway by dephosphorylation of the AKT and Forkhead box protein O1 (FoxO1)(39). In our study, we observed distinct downregulation of FoxO signaling pathway both at the system-level blood transcriptomics and during productive infection, including the FoxO transcription factors FoxO1 and FoxO3, that can act as negative feedback regulators of the innate cellular antiviral response (40). FoxO1 and FoxO3 also play an essential role in the immunometabolic dynamics and are important targets for glycolysis and gluconeogenesis (41).

One of the key pathways that were significantly upregulated both in patients’ transcriptomics and during progressive infection in Huh7 cells was OXPHOS. This indicates that CCHFV may manipulate mitochondrial dynamics for its replication by activating the OXPHOS machinery to meet elevated energy demands. Several RNA viruses like respiratory syncytial viruses (RSV), HCV, DENV, Zika virus (ZIKV) and pathogenic human coronaviruses, and are known to target mitochondria for their replication (42). Our data also showed that upon suppression of the glycolysis and glutaminolysis that fuels mitochondrial OXPHOS, there was inhibition of CCHFV-replication, further supporting the role of mitochondrial metabolism and biogenesis in CCHFV-replication and pathogenesis. Further investigations on role of mitochondrial biogenesis on CCHFV-pathogenesis can aid novel antiviral strategies.

A shift in OXPHOS can also affect the T-cells differentiation as observed in both patients’ transcriptomics and Huh7 proteomics data. In addition to innate immune responses, adaptive immune responses mediated mainly by T-cells play a critical role in the pathogenesis of viral infections. While we have observed an upregulation of IFN-related pathways in proteomics, there was a downregulation of genes belonging to Th1 and Th2 differentiation and T-cell receptor signaling pathways in the proteomics and the transcriptomics data. Th1 cells secrete IFN-γ, IL-2, TNF-α and are responsible for cell-mediated inflammatory reaction and tissue injury. Th2 cells secrete some of the cytokines including IL-10 and help B-cells for antibody production. During the acute viral infections, there is a cross-regulation for Th1 and Th2 activations primarily mediated by IL-10 and IFN-γ, respectively. Furthermore, activation of Th1 response tends to recovery from an infection while a Th2 activation results a severe clinical pathology (43). Th1 and Th2 harmonize the cell-mediated and the humoral response respectively and Th1/Th2 balance has been linked to the prognosis of viral diseases (44). Dengue hemorrhagic fever (DHF), a severe form of dengue fever (DF), is characterized by shock, hemorrhage, and death. It was shown a shift from the predominance of Th1-type response in cases of DF to the Th2-type in cases of DHF (45). Mouse model studies have shown activation of the Th1 response is associated with better protection to CCHFV-infection (46, 47). While activation of Th2 is often associated with disease severity in viral hemorrhagic fever (48), in case of CCHFV, balanced Th2-response was shown to be protective in immunized mice with a dynamic shift from Th1 to Th2 at the later part of infection (47). Both our patient data and cell infection data suggest that the virus subverts this adaptive immune response by suppressing T-cell response that could influence the disease outcome and recovery. However, this suppression did not have any impact on patient survival in our cohort through Th2 cytokines IL8 and IL10 were significantly elevated in the serum of severe cases. Our data indicated a down-regulation of Th1 and Th2 cell differentiation during acute phase of infection and at the early phase of viral replication. The naïve T cells are dependent on OXPHOS while activated T-cells on glycolysis and after differentiation, the cells are mainly dependent upon the glycolysis than OXPHOS (49). Switch in the OXPHOS during the CCHFV-infection and imbalance in Th1, Th2 and Th17 differentiation can alter the outcome of the adaptive immune response in survived CCHFV-infected patients.

One of the primary antiviral defense mechanisms is the type-I interferon (IFN-I) response. IFN-I are pleiotropic cytokines with varied cellular functions mediated by the transcriptional activations of several interferon-stimulated genes (ISGs). It is known that CCHFV-replication is sensitive to IFN-I (50). However, the virus can also delay the induction of IFN, and IFN treatment is ineffective following the establishment of infection, suggesting that CCHFV has developed mechanisms to block innate immune responses (51). The protective role of IFN-I against CCHFV has been exemplified in animal models in which IFNAR^−/−^ or STAT-1^−/−^ mice (52, 53) or STAT2^−/−^ hamsters (54) showed enhanced susceptibility to CCHFV-infection. Even in *in vitro* experiments, pre-treatment of cells with IFN-α was found to be inhibitory to CCHFV (51). Though CCHFV is inhibited by the IFN-response, not many ISGs with anti-CCHFV activity have been identified apart from MxA, although ISG20 and PKR have been proposed (55) to have anti-CCHFV activity. In the present study we observed that several ISGs with known or proposed anti-CCHFV activity, *i*.*e*., Mx1, ISG15 and ISG20 or not defined for CCHFV like IFIT1, IFIT3, IFITM3, IFI16 and OAS3 were upregulated in the acute phase CCHFV patient samples as well as in the cell-infection model.

Our CCHFV infected Huh7 proteomics data is further strengthened by a recent transcriptomics study performed in CCHFV infected Huh-7 and HepG2 showing significant alterations in IFN-response and upregulation of IFIT1, Mx1, ISG15, IF16 genes in CCHFV-infected Huh7 cells (56) as was observed in our proteomics data. The CCHFV-induced ISGs either alone or in combination with other ISGs can possess specific antiviral activities and act in regulation of IFN-signaling (57). The changes in the protein abundance of several ISGs at 24hpi and 48hpi also suggest that they have a dynamic activity during different phases of the virus infection. Furthermore, CCHFV has also evolved mechanisms to evade the immune response through the proteins they express and modifications in the genome (58, 59).

In conclusion, our study comprehensively describes the host-immune response against CCHFV that can explain viral pathogenesis. The interplay of the metabolic reprogramming towards the CCEM and its negative association with biological signaling pathways like Notch/FoxO and PI3K/mTOR/HIF-1 and the IFN-mediated host antiviral mechanism could provide attractive options for therapeutic intervention of CCHF. Further studies on the role of mitochondrial biogenesis and dynamics in CCHFV-infection, replication and pathogenesis will enhance our understanding of host-virus interactions, leading to the development of new antiviral strategies. Moreover, targeting the central carbon and energy metabolism and components of OXPHOS can be an attractive host-directed therapy during the acute CCHFV-infection by increasing the host antiviral response.

## Materials and Methods

### Ethics statement and biosafety

This study was approved by the Local Research Ethics Committee of the Ankara Numune Education and Research Hospital, Turkey (Protocol # 17-1338) and Regional Ethics Committee, Stockholm (Dnr. 2017-/1712-31/2). All patients and/or their relatives were informed about the purpose of the study and signed a consent form before collection.

### Study design, patients, and sample collection

We enrolled 18 adult patients (≥18 years) diagnosed with CCHF who were followed up by the clinical service of Infectious Diseases and Clinical Microbiology of Sivas Cumhuriyet University Hospital, Sivas, Turkey. The CCHF patients were divided into three groups using the SGS scores of 1, 2, and 3(60). Blood samples were collected on the admission day (acute stage) and from the survivors one year after their recovery (Table S1) following confirmed positive real-time RT-PCR test (Altona Diagnostics^®^, Hamburg, Germany) and/or serology by IgM indirect immunofluorescence antibody (IFA) assay (Euroimmun^®,^ Luebeck, Germany). Serum cytokine profiling targeting 22 cytokines/chemokines was performed by Public Health England using a 22 -plex customized Luminex kit (Merck Millipore, Darmstadt, Germany).

### Cells and viruses

The CCHFV strain IbAr10200 (originally isolated from *Hyalomma excavatum* ticks from Sokoto, Nigeria, in 1966) was used in this study. The small cell carcinoma in the adrenal cortex cells, SW13-ATCC^®^-CCL-105^TM,^ and human hepatocyte-derived cellular carcinoma cell line Huh7 was obtained from Marburg Virology Laboratory, (Philipps-Universität Marburg, Marburg, Germany) and matched the STR reference profile of Huh7.

### Multi-omics analysis

Peripheral blood mononuclear cells (PBMCs) RNA sequencing (RNAseq) from acute phase and convalescent phase of CCHFV-infected patients was performed as described by us recently (11, 61). Huh7 and SW13 cells were infected with CCHFV IbAr10200 at a multiplicity of infection (MOI) of 1 in triplicate, followed by tandem mass tag (TMTpro) labeled reversed-phase liquid chromatography mass-spectrometric (RPLC-MS/MS) analysis was performed as described by us recently (10, 11). The detailed methodology is presented in supplementary materials and methods.

### RNAscope and western blot

The *RNAscope*® ISH Assays (ACD Bioscience,US) targeting IFI27 (440111, ACD Bioscience, US) and CCHFV (510621, ACD Bioscience, US) were performed as described previously (62). The western blot (WB) analysis targeting RIG-I, IFIT1, ISG20, ISG15, MX1, and MX2 were performed as described by us previously (16). The WB images from all three experiments were given in Fig S6.

### Metabolic perturbation and virus infection

To inhibit glycolysis and glutaminolysis, following 1hpi (moi 0.1) the cells were treated with 2-deoxy-D-glucose (2-DG, 5mM), and diazo-5-oxo-L-norleucine (DON, 50μM) respectively. The concentrations were selected based on the minimal [mean (SD) cell viability, DON-SW13: 84% (4%), DON-Huh7: 78% (2%) and 2-DG-SW13: 80% (2%) or no cytotoxicity (2-DG in Huh7) in the respective cells 24hrs following drug treatment. The cells were collected after 24hpi and the cells were lysed in Trizol reagent. RNA was extracted using the Direct-zol RNA Miniprep kit (Zymo Research, Irvine, CA) according to the manufacturer’s instructions. Viral RNA was measured by quantitative real real-time polymerase chain reaction (qRT-PCR) using TaqMan Fast Virus 1-Step Master Mix (Thermo Fisher Scientific) with primers and probe specific for the CCHFV L gene.

### Data and code availability

All the codes are available in GitHub (https://neogilab.github.io/CCHF-Turkey/). Raw RNAseq data is available in Sequence Read Archive (SRA) with id PRJNA680886. The mass spectrometry proteomics data have been deposited to the ProteomeXchange Consortium via the PRIDE partner repository with the dataset identifier PXD022672.

## Acknowledgment

The study is funded by Swedish Research Council Grants 2018-05766 and 2017-03126 to AM and 2017-01330 to UN, PHE Grant In Aid 109509 (inc. PhD studentship [EK]) and EU-H2020 CCHFVaccine to RH. The authors would like to acknowledge support from the National Genomics Infrastructure (NGI), Science for Life Laboratory, for RNAseq and Proteomics Biomedicum; and Karolinska Institute, Solna, for LC-MS/MS analysis. Authors would like to acknowledge Dr. Shubha. Krishnan for critical reading of the manuscript and comments and Xi Chen for helping to extract RNA. The computations were performed using resources provided by SNIC through the Uppsala Multidisciplinary Center for Advanced Computational Science (UPPMAX) under Project SNIC2017-550. The microscopy part of the study was performed at the Live Cell Imaging Facility and Biomedicum Imaging Core, Karolinska Institute, Sweden, supported by grants from the Knut and Alice Wallenberg Foundation, the Swedish Research Council, the Centre for Innovative Medicine, and the Jonasson Center at the Royal Institute of Technology, Sweden.

## Supplementary Materials

### Supplementary Materials and Methods

**Fig. S1: Severity group association with gene expression. (A)** MA-plot of differentially regulated genes during the acute phase between samples of severity group 1 and severity group 2 and 3. **(B)** Sample distribution during the acute phase of infection in different severity groups as reported.

**Fig S2**. Violin plot of 22 soluble markers as determined from Luminex assay assays.

**Fig S3**. Gene set enrichment analysis of the individual communities.

**Fig S4**. Weight co-expression network of the negatively co-related genes We observed a high number of negative correlations between this community (c1) and those associated with Notch, mTOR and FoxO signaling (c5) and HIF-1 signaling (c7).

**Fig S5**. Quantitative proteomics of the Huh7 with 4 MOI infection 24hpi and comparisons with the 1 MOI infection indicated 2452 proteins were common that were significantly dysregulated. Protein set enrichment analysis identified 33 pathways were dysregulated where the top pathways remain unchanged.

**Fig S6**. Western blot Images of ISGs (RIG-I, IFIT1, Mx1, Mx2, ISG20, ISG15), CCHFV-N protein and β-actin at 48hpi from three experimental replicates.

**Table S1**. The CCHF patient characteristics.

**Dataset S1:** The DGE profile for the acute phase compared to the recovered phase in all patients.

**Dataset S2:** Pathways found to be significantly regulated by genes expressed at the acute infection phase compared to recovered phase identified in PIANO.

**Dataset S3:** Pathways found to be significantly regulated by proteins in mock and CCHFV-treated Huh7 cells following 24hpi and 48hpi and time-series analysis identified in PIANO.

