## Supplemental Figure S1 and S6 for "Multi-omics insights into host-viral response and pathogenesis in Crimean-Congo Hemorrhagic Fever Viruses for novel therapeutic target"

**This PDF file includes:**

Supplementary text
Figures S1 to S6
Tables S1
Legends for Datasets S1 to Sx
SI References

**Other supplementary materials for this manuscript include the following:**

Datasets S1 to Sx

**Materials and Methods:**

**RNA extraction and sequencing (Illumina RNAseq) and transcriptomics analysis.** Total RNA was extracted from Trizol-treated PBMC using the Direct-zol RNA Miniprep (Zymo Research, CA, USA) according to the manufacturer's protocol. RNA-Seq was performed at the National Genomics Infrastructure, Science for Life Laboratory, Stockholm, Sweden, as described by us previously (1). The transcriptomics data analysis was performed as described by us recently (2). The distribution of all samples was then visualized after reducing the dimension of the data by applying the Uniform Manifold Approximation and Projection for Dimension Reduction (UMAP) technique using R package umapv0.2.6.0. The reduced dimensions of the data were plotted in 2D space using R package ggplot2 v3.3.2 (https://cran.r-project.org/web/packages/ggplot2/index.html). Differential gene expression analysis was performed using raw read counts of remaining samples using the R/Bioconductor package DESeq2 v1.26.0 (.https://bioconductor.org/packages/release/bioc/html/DESeq2.html) Genes with adjusted p-values<0.05 were considered as significantly regulated. KEGG gene set enrichment analysis for differentially regulated genes was performed using PIANO v2.2.0(3) (nperm=500, geneset statistic =mean) and for communities using enrichr module of python package GSEAPY v0.9.16 (https://github.com/zqfang/GSEApy). Pathways belong to KEGG category of metabolism, environmental information processing and organismal systems were used for the analysis. Additionally, three gene sets related to IFN-signaling curated by the group (4) were also considered for the enrichment analysis of gene communities. GO enrichment analysis was performed using enrichr for GO biological process 2018 (https://maayanlab.cloud/Enrichr/). Redundant GO terms were removed using the online tool REVIGO (5). Reporter metabolites(6) were identified through PIANO using the genome-scale metabolic model Human1 (7) to generate the metabolite-gene sets. Heatmaps were generated using the R/Bioconductor package ComplexHeatmapv2.2.0 (https://www.bioconductor.org/packages/release/bioc/ html/ComplexHeatmap.html.) Bubble plots, MA plots, volcano plots, violin plots and bar plots were created using R package ggplot2 v3.3.2. Network visualization was performed using Cytoscape v3.6.1 (https://cytoscape.org/). Networks were built by computing pairwise Spearman rank correlations between all genes after removal of non-expressed (row median FPKM<1) or lowly variant (row variance < 0.1) genes, and analyzed in igraph for those displaying statistically significant (adjusted P<0.001 positive correlations. Centrality analysis was performed by computing degree centrality. Communities were identified by modularity maximization through the Leiden algorithm (8). Venn diagrams were constructed using the online tool InteractiVenn (http://www.interactivenn.net/). All the codes are available in GitHub (https://neogilab.github.io/CCHF-Turkey/). Raw RNAseq data is available in Sequence Read Archive (SRA) with id PRJNA680886.

***In vitro* infection assay in Huh7 cells.** Huh7 and SW13 cells were infected with the CCHFV in triplicate, as described by us previously (2). Briefly, Huh7 cells were infected with CCHFV IbAr10200 at a multiplicity of infection (MOI) of 1. After 1 h of incubation (37°C, 5% $CO_2$) the inoculum was removed, the cells were washed with PBS, and 2 ml DMEM supplemented with 5% heat-inactivated FBS was added to each well. Samples were collected in triplicate at 24 and 48 hpi along with controls. Due to high permissivemess we restricted the SW13 infection of 1 MOI for 24hrs only. The infection in Huh7 24hpi were confirmed by immunofluorescence staining of CCHFV nucleoprotein-protein (Fig S10). The cells were fixed in ice-cold acetone-methanol (1:1) and stained using a rabbit polyclonal anti-CCHFV nucleocapsid antibody (home-made) followed by a fluorescein isothiocyanate (FITC)-conjugated anti-rabbit antibody (Thermo Fisher Scientific, US) and DAPI (Roche, US).

**Tandem mass tag (TMTpro) labelled reversed-phase liquid chromatography mass-spectrometric (RPLC-MS/MS) analysis**. The RPLC-MS/MS of the TMTpro labelled samples was performed as described by us recently (2, 4). Briefly, following the protein digestion in S-Trap microcolumns (Protifi, Huntington, NY), the resulting peptides were labeled with TMTpro tags. Labeled peptides were fractionated by high pH (HpH) reversed-phase chromatography, and each fraction was analyzed on an Ultimate 3000 UHPLC (Thermo Scientific, San Jose, CA) in a 120 min linear gradient. Proteins were searched against the SwissProt human database using the search engine Mascot v2.5.1 (MatrixScience Ltd, UK) in Proteome Discoverer v2.4 (Thermo Scientific) software allowing up to two missed cleavages.

**Proteomics data analysis.** The raw data were first filtered to remove missing data. Proteins detected in all samples were retained for analysis resulting in 8,501 proteins in the filtered dataset. The filtered data was then normalized by applying eight different methods using R/Bioconductor package NormalyzerDE v1.4.0 (http://quantitativeproteomics.org/normalyzerde). The quantile normalization was found superior to other methods and was selected for further use. Differential protein expression analysis was performed using R/Bioconductor package limma v3.42.2 (https://bioconductor.org/packages/release/bioc/html/limma.html). Proteins with adjusted p-values of less than 0.05 were regarded as significant. KEGG pathway enrichment analysis was performed as mentioned in the transcriptomics section. The mass spectrometry proteomics data have been deposited to the ProteomeXchange Consortium via the PRIDE partner repository with the dataset identifier PXD022672.

**RNAscope targeting IFI27 (ISG12) and CCHF.** The *RNAscope*® ISH Assays (ACD Bioscience,US) targeting IFI27 (440111, ACD Bioscience, US) and CCHFV (510621, ACD Bioscience, US) were performed as described previously(9). SW13 cells were infected with CCHFV (Ibar 10200 strain) at MOI of 0.1. After 1h, inoculum was replenished with fresh Leitbovitz medium containing 5% FBS and incubated 48h. After infection cells were fixed 30 min using ice-cold acetone. Cells were permeabilized using 0.1% Triton x-100 for 10 min at RT prior probe incubation at 40□C of IFI27 and CCHFV probes (RNAscope® ISH ACD, US) for 2h, 30min, 15 min, 30 min and 15 min, respectively. Cells were further stained using DAPI and coverslip attached using ProLong Gold Antifade reagent (P10144, Thermofisher). Images were acquired using Nikon Single Point scanning confocal with 60/1.4 oil objective.

**Metabolic perturbation and virus infection.** To inhibit glycolysis and glutaminolysis, following 1hpi (moi 0.1) the cells were treated with 2-deoxy-D-glucose (2-DG, 5mM), and diazo-5-oxo-L-norleucine (DON, 50□M)) respectively. The concentrations were selected based on the minimal [mean (SD) cell viability, DON-SW13: 84% (4%), DON-Huh7: 78% (2%) and 2-DG-SW13: 80% (2%) or no cytotoxicity (2-DG in Huh7) in the respective cells 24hrs following drug treatment. The cells were collected after 24hpi and the cells were lysed in Trizol reagent. RNA was extracted using the Direct-zol RNA Miniprep kit (Zymo Research, Irvine, CA) according to the manufacturer's instructions. Viral RNA was measured by quantitative real real-time polymerase chain reaction (qRT-PCR) using TaqMan Fast Virus 1-Step Master Mix (Thermo Fisher Scientific) with primers and probe specific for the CCHFV L gene; Forward: 5-GCCAACTGTGACKGTKTTCTAYATGCT-3', Reverse-1: 5'- CGGAAAGCCTATAAAACCTACCTTC-3', Reverse-2: 5'-CGGAAAGCCTATAAAACCTGCCYTC-3' and Reverse-3: 5'-CGGAAAGCCTAAAAAATCTGCCTTC-3' and Probe FAM-CTGACAAGYTCAGCAAC –MGB. RNAse was used as endogenous control. The cycling reactions was performed using a capillary Roche LightCycler 2.0 system.

**Western Blot.** Huh7 cells were infected with CCHFV at MOI of 1 and 48hpi cells were harvested and lysed in 10% SDS lysis buffer (10 mM Tris-Cl, 150 mM NaCl, 10% SDS, freshly supplemented with 1× protease inhibitor cocktail) to which NuPAGE™ LDS sample buffer (ThermoFisher Scientific,US) was added and the samples were boiled at 99°C for 10 min. The protein concentration was evaluated by Pierce™ 660nm Protein Assay kit (ThermoFisher Scientific, US). Evaluation of protein expression was performed by running 20µg of total protein lysate on NuPage Bis Tris 4%–12% (Invitrogen, Carlsbad, CA, USA). Proteins were transferred using iBlot dry transfer system (Invitrogen, Carlsbad, CA, USA) and blocked for two hours using 5% milk in Tris-buffered saline containing 0.1% Tween-20(TBSt). Subsequent antibody incubation was performed at 4°C overnight. Membranes were washed using 0.1% TBSt and secondary antibody was incubated for one and half hour at room temperature using Dako polyconal goat anti-rabbit or anti-mouse Immunoglobulins/HRP (Aglient Technologies, Santa Clara, CA, USA). Membranes were washed using 0.1% TBSt and proteins were detected using ECL or ECL Select (GE Healthcare, Chicago, IL, USA) on ChemiDoc XRS+ System (Bio-Rad Laboratories, Hercules, CA, USA). The western blot analysis was performed by using the antibodies presented in Table S4.

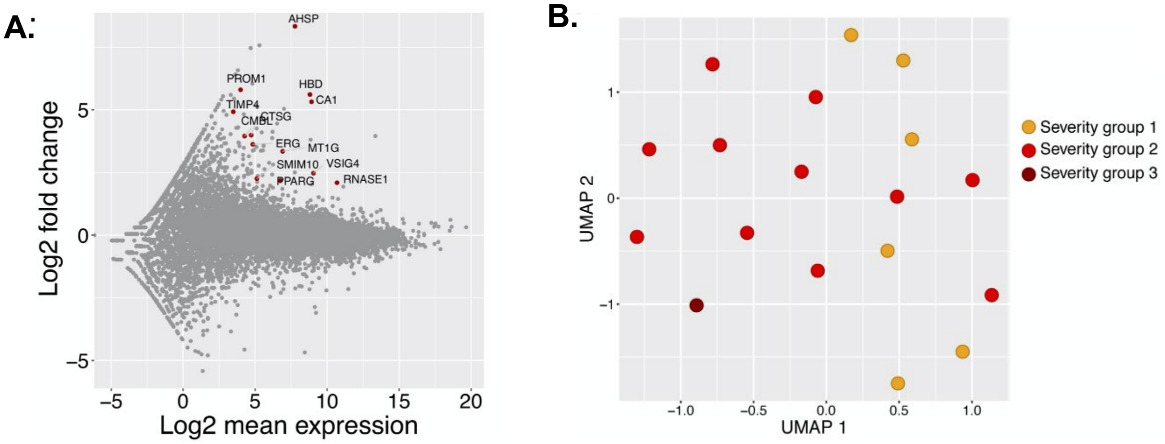


**Fig. S1: Severity group association with gene expression. (A)** MA-plot of differentially regulated genes during the acute phase between samples of severity group 1 and severity group 2 and 3. **(B)** Sample distribution during the acute phase of infection in different severity groups as reported.

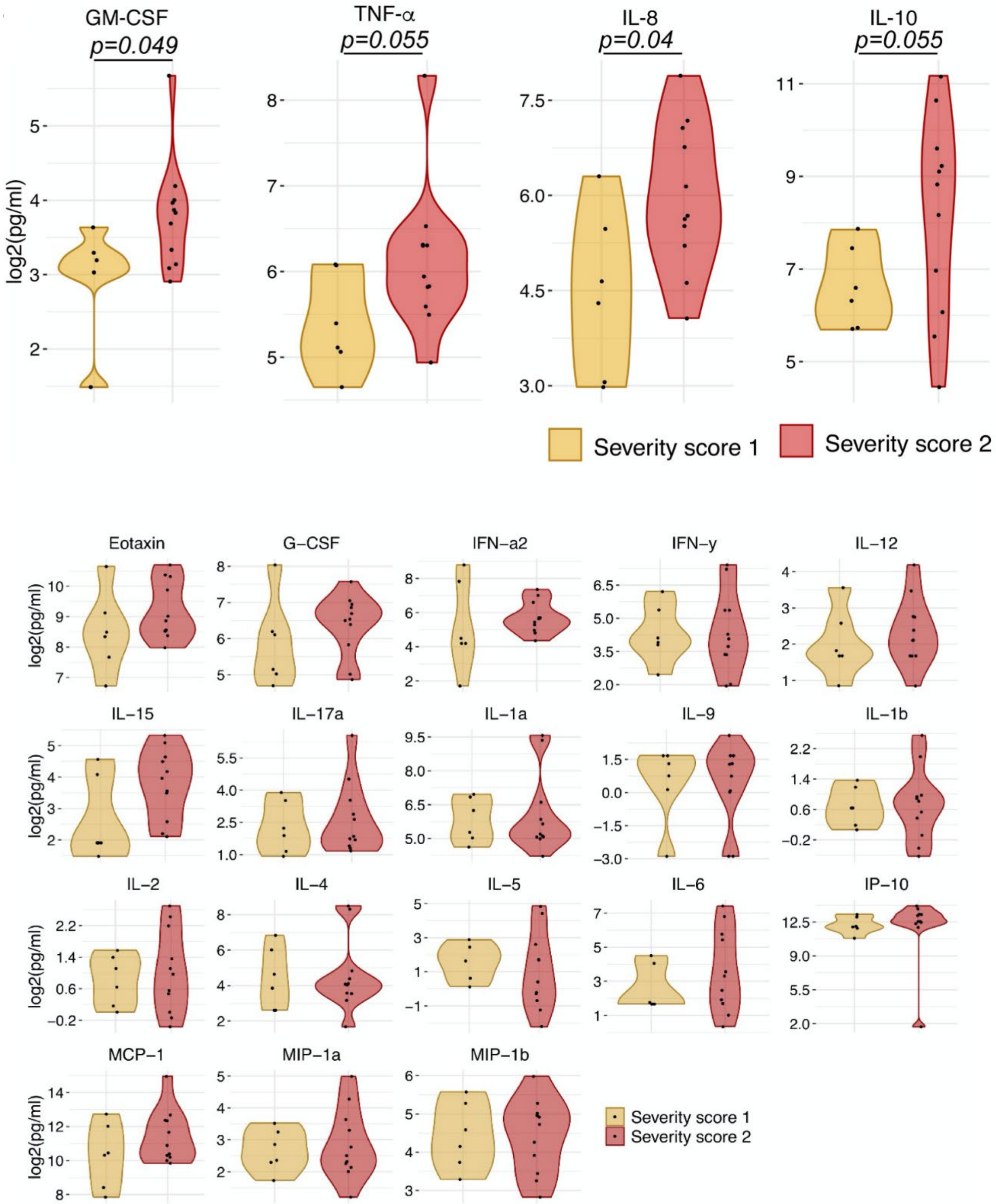


**Fig S2.** Violin plot of 22 soluble markers as determined from Luminex assay assays.

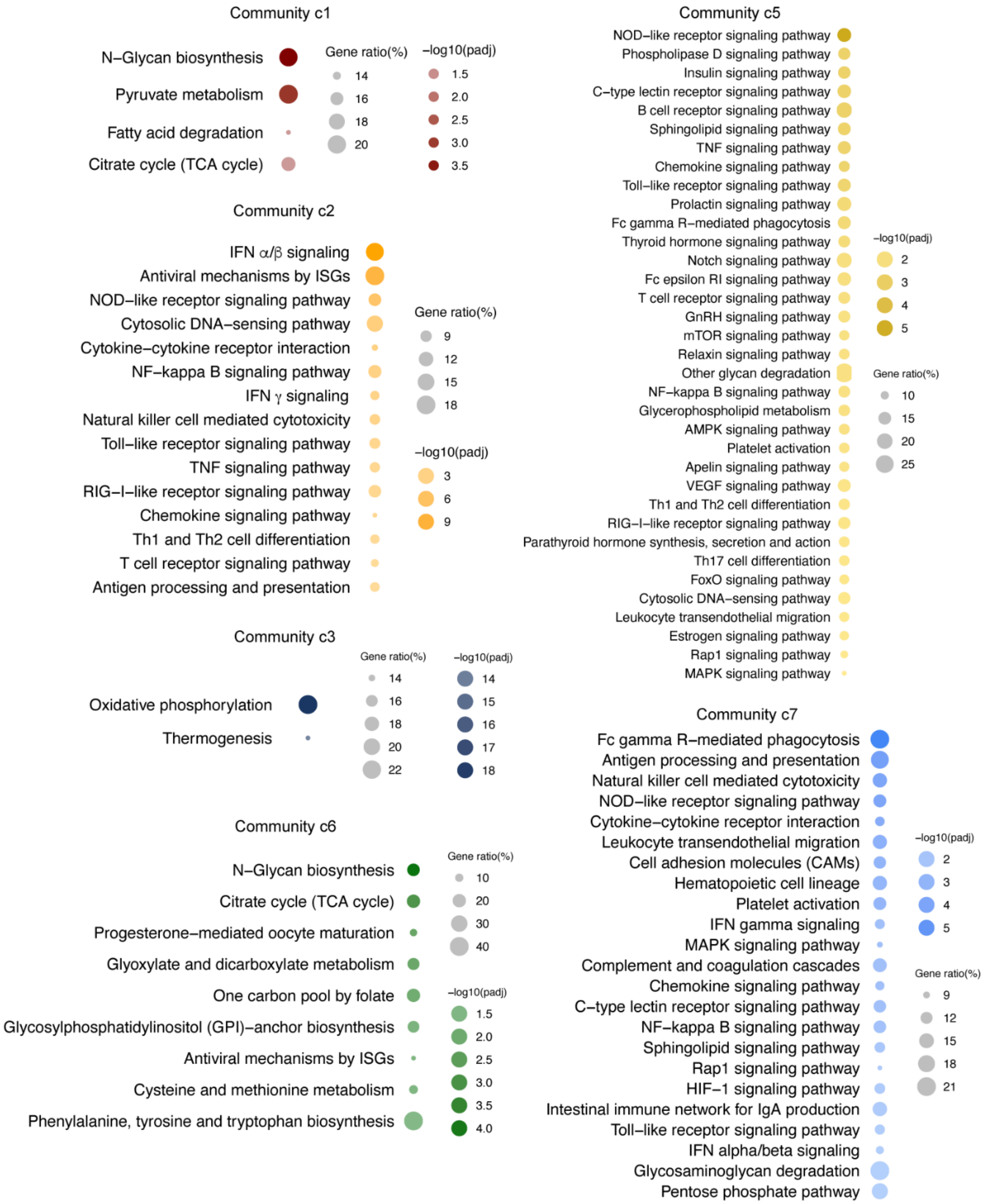


**Fig S3**. Gene set enrichment analysis of the individual communities.

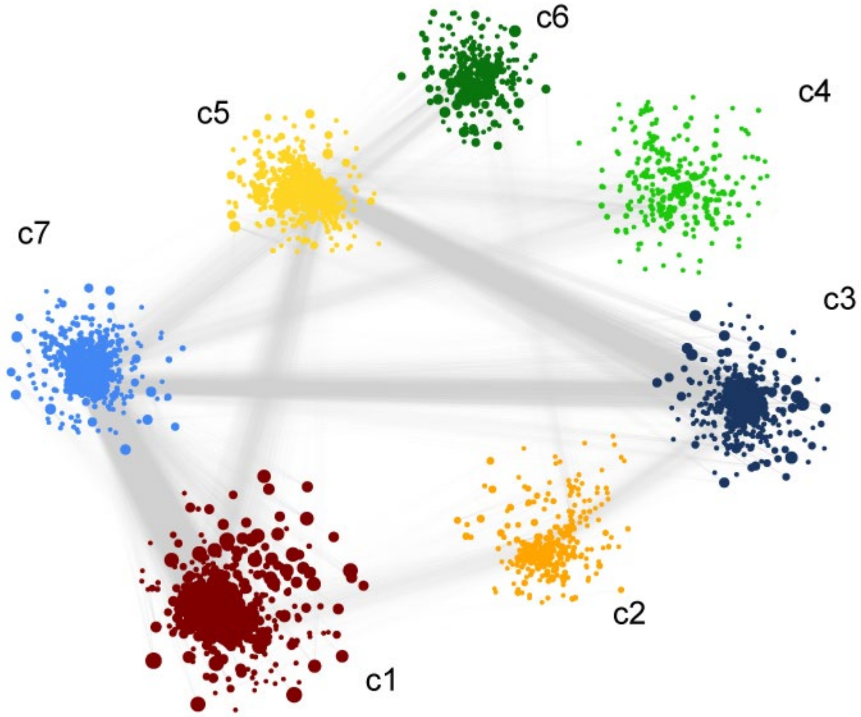


**Fig S4**. Weight co-expression network of the negatively co-related genes We observed a high number of negative correlations between this community (c1) and those associated with Notch, mTOR and FoxO signaling (c5) and HIF-1 signaling (c7).

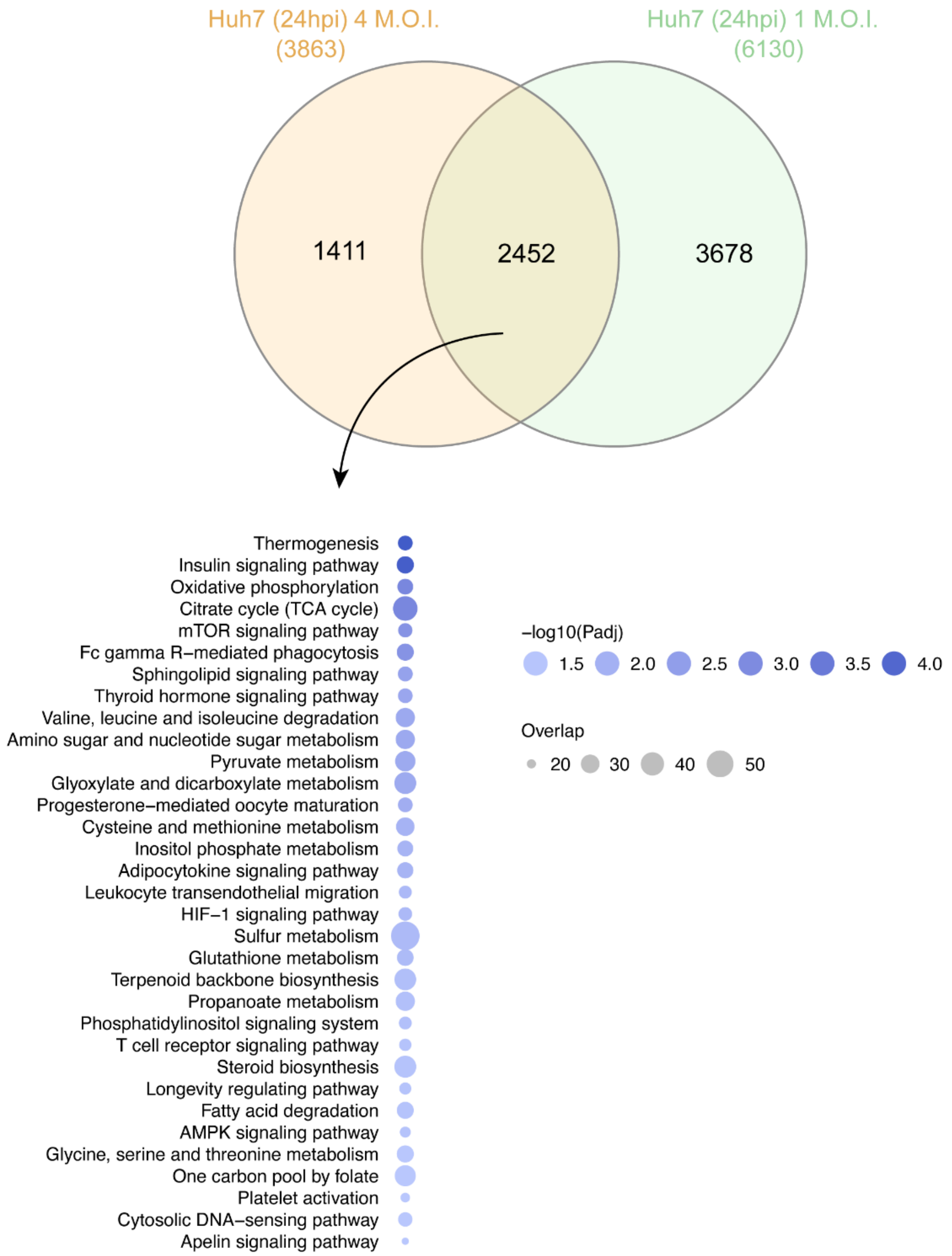


**Fig S5.** Quantitative proteomics of the Huh7 with 4 MOI infection 24hpi and comparisons with the 1 MOI infection indicated 2452 proteins were common that were significantly dysregulated. Protein set enrichment analysis identified 33 pathways were dysregulated where the top pathways remain unchanged.

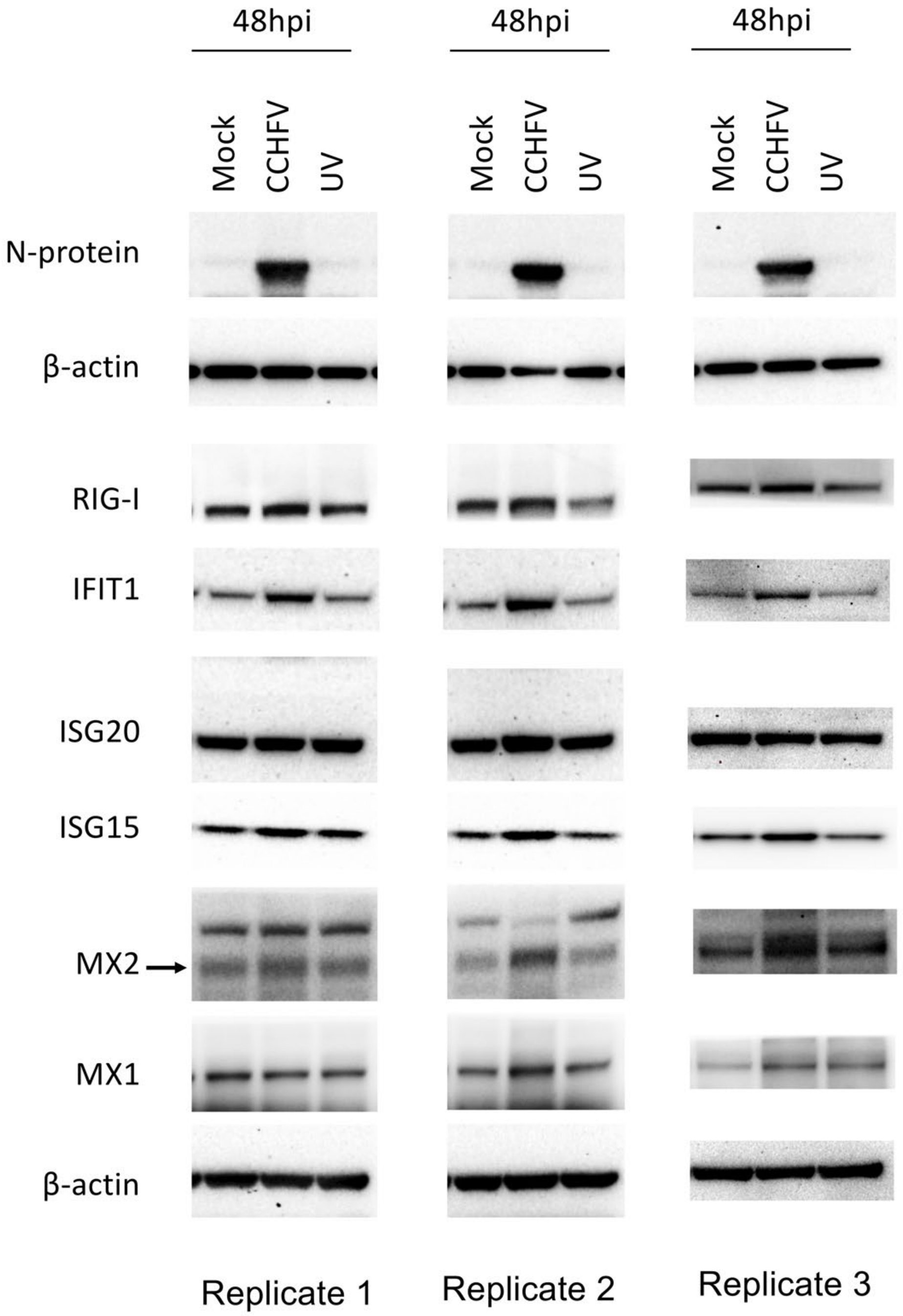


**Fig S6.** Western blot Images of ISGs (RIG-I, IFIT1, Mx1, Mx2, ISG20, ISG15), CCHFV-N protein and β-actin at 48hpi from three experimental replicates.

**Table S1.** The CCHF patient characteristics.

| PID | Age | Gender | The date of symptoms onset | The date of hospitalization | Time to hospitalization (days) | The date of the first sampling | The date of the second sampling | SGS score | Severity group* | RT-PCR | CT-values | Anti-CCHFV IgM | Outcome |
|---|---|---|---|---|---|---|---|---|---|---|---|---|---|
| **P01** | 33 | Female | 30 May 2017 | 03 June 2017 | 4 | 03 June 2017 | 05 July 2018 | 5 | 1 | positive | 31,85 | ND | Survived |
| **P02** | 18 | Male | 06 June 2017 | 12 June 2017 | 6 | 12 June 2017 | ND | 7 | 2 | positive | 25,89 | positive | Survived |
| **P03** | 45 | Male | 12 June 2017 | 13 June 2017 | 1 | 14 June 2017 | 01 July 2018 | 0 | 1 | positive | 21,87 | ND | Survived |
| **P04** | 67 | Male | 13 June 2017 | 16 June 2017 | 3 | 17 June 2017 | 05 July 2018 | 8 | 2 | positive | 22,38 | ND | Survived |
| **P05** | 48 | Male | 12 June 2017 | 18 June 2017 | 6 | 19 June 2017 | 08 July 2018 | 7 | 2 | positive | 29,79 | ND | Survived |
| **P06** | 68 | Male | 13 June 2017 | 19 June 2017 | 6 | 20 June 2017 | 05 July 2018 | 5 | 1 | positive | 28,41 | ND | Survived |
| **P07** | 77 | Male | 19 June 2017 | 22 June 2017 | 3 | 23 June 2017 | 05 July 2018 | 6 | 2 | positive | 24,77 | ND | Survived |
| **P08** | 29 | Female | 20 June 2017 | 24 June 2017 | 4 | 25 June 2017 | 02 July 2018 | 6 | 2 | positive | 26,91 | ND | Survived |
| **P09** | 50 | Female | 20 June 2017 | 25 June 2017 | 5 | 26 June 2017 | 06 July 2018 | 4 | 1 | positive | 26,36 | positive | Survived |
| **P10** | 35 | Female | 07 July 2017 | 12 July 2017 | 5 | 12 July 2017 | 04 July 2018 | 3 | 1 | negative | NA | positive | Survived |
| **P11** | 64 | Female | 15 July 2017 | 18 July 2017 | 3 | 19 July 2017 | ND | 10 | 2 | positive | 22,46 | ND | Survived |
| **P12** | 57 | Male | 16 July 2017 | 21 July 2017 | 5 | 22 July 2017 | 09 July 2018 | 9 | 2 | positive | 20,81 | ND | Survived |
| **P13** | 79 | Male | 22 July 2017 | 24 July 2017 | 2 | 24 July 2017 | ND | 11 | 3 | positive | 22 | ND | Died |
| **P14** | 36 | Male | 01 August 2017 | 06 August 2017 | 5 | 07 August 2017 | 04 July 2018 | 7 | 2 | positive | 24,66 | ND | Survived |
| **P15** | 62 | Male | 15 August 2017 | 20 August 2017 | 5 | 21 August 2017 | 06 July 2018 | 9 | 2 | positive | 19,86 | ND | Survived |
| **P16** | 48 | Male | 05 September 2017 | 07 September 2017 | 2 | 07 September 2017 | ND | 4 | 1 | positive | 22,09 | ND | Survived |
| **P17** | 55 | Male | 12 April 2018 | 17 April 2018 | 5 | 18 April 2018 | ND | 9 | 2 | positive | 26,16 | ND | Survived |
| **P18** | 44 | Female | 23 April 2018 | 27 April 2018 | 4 | 29 April 2018 | ND | 9 | 2 | positive | 21,27 | ND | Survived |

* 1: Low (0-5); 2: Intermediate (6-10); 3: High (11-16)

ND: not determined; NA: not applicable; SGS: severity grading system; RT-PCR: real time - polymerase chain reaction; CT: cycle threshold; CCHFV: Crimean-Congo haemorrhagic fever virus

**Dataset S1:** The DGE profile for the acute phase compared to the recovered phase in all patients.

**Dataset S2:** Pathways found to be significantly regulated by genes expressed at the acute infection phase compared to recovered phase identified in PIANO.

**Dataset S3:** Pathways found to be significantly regulated by proteins in mock and CCHFV-treated Huh7 cells following 24hpi and 48hpi and time-series analysis identified in PIANO.
